# Ends and middle: global force balance determines septum location in fission yeast

**DOI:** 10.1101/520007

**Authors:** Xavier Le Goff, Jordi Comelles, Charles Kervrann, Daniel Riveline

## Abstract

The fission yeast cell is shaped as a very regular cylinder ending by hemi-spheres at both cell ends. Its conserved phenotypes are often used as read-outs for classifying interacting genes and protein networks. Using Pascal and Young-Laplace laws, we proposed a framework where scaling arguments predicted shapes. Here we probed quantitatively one of these relations which predicts that the division site would be located closer to the cell end with the larger radius of curvature. By combining genetics and quantitative imaging, we tested experimentally whether altered shapes of cell end correlate with a displaced division site, leading to asymmetric cell division. Our results show that the division site position depends on the radii of curvatures of both ends. This new geometrical mechanism for the proper division plane positioning could be essential to achieve even partitioning of cellular material at each cell division.

## 1. Introduction

Biological screens are often based on the classification of shared phenotypes. This approach is used successfully for a variety of model systems, ranging from yeast [1] to *C. elegans* [2], Zebrafish [3] and *Drosophila* [4]. In fission yeast, for example, this approach has allowed to reveal essential genes involved in a large class of phenomena such as cell shape [5,6], polarity [7], cell fusion [8], cell cycle [9], nuclear volume [10], and septum position [11]. Interacting proteins can be labeled fluorescently and their localisations in mutants mapped over time to correlate placements and distribution of proteins with potential activating and inhibiting effects [12–15].

However, these genetic changes in strains are often associated also with changes in cell shapes. With this respect, differences in pressure between the inside and the outside of the cell together with a constant surface tension can be utilised to derive simple laws for read-outs, such as localisation of polarity cues for lower tension, displacements of division planes, buckled mutants [16]. This approach has proven its potential impact for new rules of self-organisations and localisations [17].

We proposed in reference [16] scaling relations for shapes using Pascal principle and Young-Laplace law. Here we test one of these relations quantitatively using fission yeast, the placement of the septum. Its biological function is essential, since this physical separation between sister cells secures the reliable partition of biological materials at the end of each cell cycle.

The cell is shaped as a very regular cylinder ending by hemi-spheres at both cell ends. Fission yeast cells elongate during interphase keeping this regular shape, set by a balance between cell wall stiffness and turgor pressure [16]. The cell is being remodelled locally at the cell ends to promote cell extension. The nucleus is permanently maintained in the middle of the cell by different forces, including microtubule pushing forces [18,19]. The central position of the nucleus is used as a spatial cue to assemble a contractile actomyosin ring when cells entered mitosis. This ring is used to drive the synthesis of a specific cell wall structure called the division septum, which physically separates the daughter cells into two cells and leaving them approximately half the mother cell after cytokinesis. We proposed scaling arguments where the radii of curvatures of cell ends *R_i_* influence constraints on the cell wall [16]. They ultimately produce an effect on the division site position. This relation predicts that a higher radius of curvature at one cell end should displace the division site by a length *L_shift_* due to unbalanced stress applied on the cell wall. The division site would be located closer to the cell end with the larger radius (lower curvature, defined as 1/R). We decided to test experimentally whether altered shapes of cell end actually correlate with a displaced division site, leading to an asymmetric cell division. We combined genetics together with live cell imaging. We used two strains modified from the wild type strain, *i.e*. a constitutive deletion *tea4Δ* mutant and a conditional *kin1-as1* mutant, which affect cell ends. Our results show that the division site position depends on the radii of curvatures of both ends.

## 2. Experimental procedures

### 2.1. Yeast Strains and General Techniques

*S. pombe* strains used in this study are XLG52 (*h- cdc15::GFP-ura4^+^ ura4-D18 leu1-32*, a kind gift of V. Simanis, Switzerland), XLG540 (*h- tea4::ura4+ ura4-D18 cdc15::GFP-ura4^+^ leu1-32*), XLG741 (*h- cdc15::GFP-ura4^+^ kin1-as1 leu1-32 ura4-D18*). Growth media and basic techniques for *S. pombe* have been described [20].

### 2.2. Microscopy

A spinning disk Nikon TE2000 microscope, equipped with a 100x 1.45 NA PlanApo oil-immersion objective lens and a HQ2 Roper camera, was used for data acquisition. Cells expressed the acto-myosin ring component Cdc15-GFP and were stained 10 minutes with isolectin-488 (Molecular probes) that stains the global cell wall (but not the septum). Metamorph software was used for capturing images. The “three point circular” ImageJ Plugin allows to draw a ring with three points at a cell end and it gives the radius of curvature. We used this Plugin to measure radii of curvature to obtain the best measurements. Cell lengths (L, L1 and L2) were measured with the Plot profile Plugin. For Transmission Electron Microscopy, cells were stained with potassium permanganate and images were captured by a Jeol Jem-1010 (Peabody, MA).

### 2.3. Statistical analyses and graphical representation

Statistical analysis was done using GraphPad Prism. Results are presented as mean ± s.e.m of N = 3 experiments, n_WT_ = 52 cells, n_*tea4Δ*_ = 51 cells, n_*kin1-as1*+DMSO_ = 35 cells and n_*kin1-as1*+NMPP1_ = 48 cells. First, normality of the datasets was tested by the d’Agostino-Pearson normality test. Statistical differences were analyzed by t-test (Gaussian distribution) and Mann-Whitney test (non Gaussian distribution). The Pearson’s r correlation coefficient (Gaussian distribution) and the Spearman correlation coefficient (non Gaussian distribution) were used in order to test the relation between (R_1_-R_2_) and L_shift_ for all the conditions. (R_1_-R_2_) as function of L_shift_ were fitted using a linear regression. To obtain the mean surface tension *γ* for *tea4*Δ and *kin1-as1* cells, *γ* was taken constant *on average* at cell ends, and calculated by rewriting Equation 10 from reference [16] to give *l_shift_* = *(<γ_end_ >/(ΔP·R_side_))·(R_1_-R_2_)* (Eq. 1), where L_shift_, R, R_1_ and R_2_ are obtained from the experiments and assuming ΔP = 0.85 MPa for every cell. The distribution of <*γ_end_*> was plotted and fitted with a Gaussian distribution y = A·exp(-(x-x_c_)^2^/(2·w^2^)), where x_c_ is the mean value for *γ_end_* and w corresponds to the standard deviation.

## 3. Results

According to our scaling law, the cell end curvature would impact on cell division site position at the time of septum ingression due to cell wall constraints. Therefore, we monitored the cell division site localization using the expression of the cytokinetic ring component Cdc15-GFP (Figure 1A). Cellular outlines were stained with the cell wall isolectin-488 label. Equation 10 of reference [16] showed that this shift with actual center does not depend on the longitudinal cell length *L* (Figure 1B) and can be rewritten as *l_shift_* = *(<γ_end_>/(ΔP·R_side_))·(R_1_-R_2_*), (Eq. 1), where *R* is the mean radius of the long axis, *ΔP* the constant difference in pressure between the inside and the outside of the cell, if we consider the mean cell wall surface tension *<γ_end_>*.

**Figure 1.**
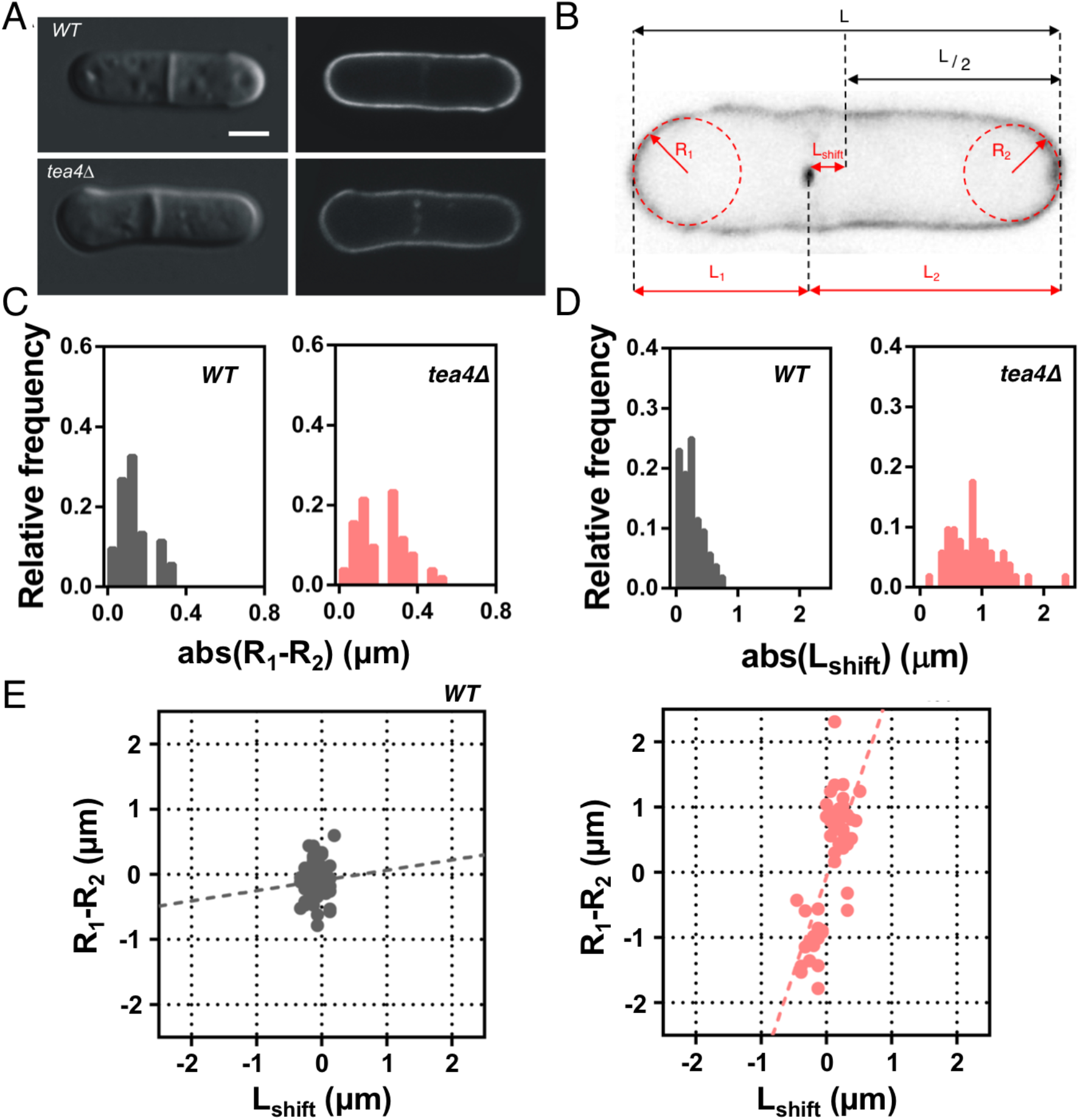
**(A)** DIC (left) and fluorescent (right) microscopy images (cdc15-GFP, isolectin-488) of WT and tea4D, Scale bar 2μm. **(B)** Schematics of the parameters measured in each cell: R_i_ corresponds to the radii of curvature at the cell “end i “, and L_i_ corresponds to the distance between “end i “ and septum; L_shift_ is defined as the distance between the septum and the middle plane (L_shift_=(L_2_-L_1_)/2), L the total length. **(C)** Distributions of the absolute values of R_1_-R_2_ for the WT and tea4D cells. Ro-R; distributions are statistically different (t-test, p = 0.0003). **(D)** Distributions of the absolute values of L_shift_ values for the WT and tea4D cells. Distributions are statistically different (Mann-Whitney, p < 0.0001). **(E)** Correlation between L_shift_ and R_1_-R_2_ for the WT and tea4D cells. The fits of Equation 1 are shown in the graphs. n_WT_=52 and n_tea4Δ_=51. Spearman correlation coefficients are 0.03 and 0.61 respectively.

First, we compared wild type (WT) and the asymmetrically dividing mutant *tea4Δ* (Figure 1A). The Tea4 protein is involved in bipolar activation of cell growth in the cell ends. *tea4Δ* cells showed altered cell morphology including one more enlarged cell end than the other and an asymmetrically positioned division site [21,22], that was determined with Cdc15-GFP. We calculated *L_shift_* value as *L_shift_*=*(L_2_-L_1_)/2* (Eq. 2), where L_1_ stands for the distance between the ring and one cell end (cell end1) and L_2_ stands for the distance between the ring and the cell end (cell end2). To calculate the *R_1_-R_2_* value, radii of curvatures were measured as follows: *R_1_* for cell end1 (or the end associated to L_1_) and *R_2_* for the other cell end (or the end associated to L_2_). We observed an increased asymmetry in *tea4Δ* cells compared to WT, both for R_1_-R_2_ value (Figure 1C) and L_shift_ value (Figure 1D). On one hand, cell end radii are different in *tea4Δ* cells compared to WT cells. Difference of radii of curvature at the cell ends (*R_1_-R_2_*) augmented from 0.13±0.01 μm for WT cells to 0.21±0.02 μm for *tea4Δ* (Figure 1C), showing an increase of the asymmetry in the cap curvatures. Although a small asymmetry for WT cells was observed, probably due to the intrinsic noise of the biological system, it significantly changed about two-fold for *tea4Δ* mutant. On the other hand, the mean value of *L_shift_* increased from 0.26±0.05 μm for WT cells to 0.62±0.06 μm for *tea4Δ* cells (Figure 1D). Again, a non-zero *L_shift_* was observed for WT cells attributed to the intrinsic variability of biological systems. However, the _shift_ in the division site significantly increased by two-fold in *tea4Δ* cells. Thus, there is clearly a larger amplitude of *L_shift_* in *tea4Δ* cells indicating that they divide more asymmetrically than WT cells. Finally, the *L_shift_*, plotted as a function of the *R_1_-R_2_* difference of radii of cell end curvatures (Figure 1E), showed a positive correlation between *L_shift_* and *R_1_-R_2_* for *tea4Δ* cells (correlation coefficient 0.61), whereas no correlation was observed for WT cells (correlation coefficient 0.03). Therefore, the experimental results show that the division site is displaced towards the end with the highest radius of curvature, which is consistent with our prediction.

*Tea4Δ* cells are constitutively misshapen and cell end curvatures differences may arise from cell wall defects inherited through several generations independently of cell division site selection. Thus, we used *kin1-as1*, a conditional allele of the cell wall regulating Kin1 kinase, that promoted cell division site mispositioning within the duration of a cell division cycle. *Kin1-as1* was inhibited using a small molecule called NMPP1 added into the culture medium [23]. *Kin1-as1* NMPP1 and *kin1-as1* DMSO cells are isogenic but cultured with or without the inhibitor for 2 hours, respectively. *Kin1-as1* NMPP1 cells adopt an asymmetric cell division pattern in less than a generation time (see Figure 2A).

**Figure 2.**
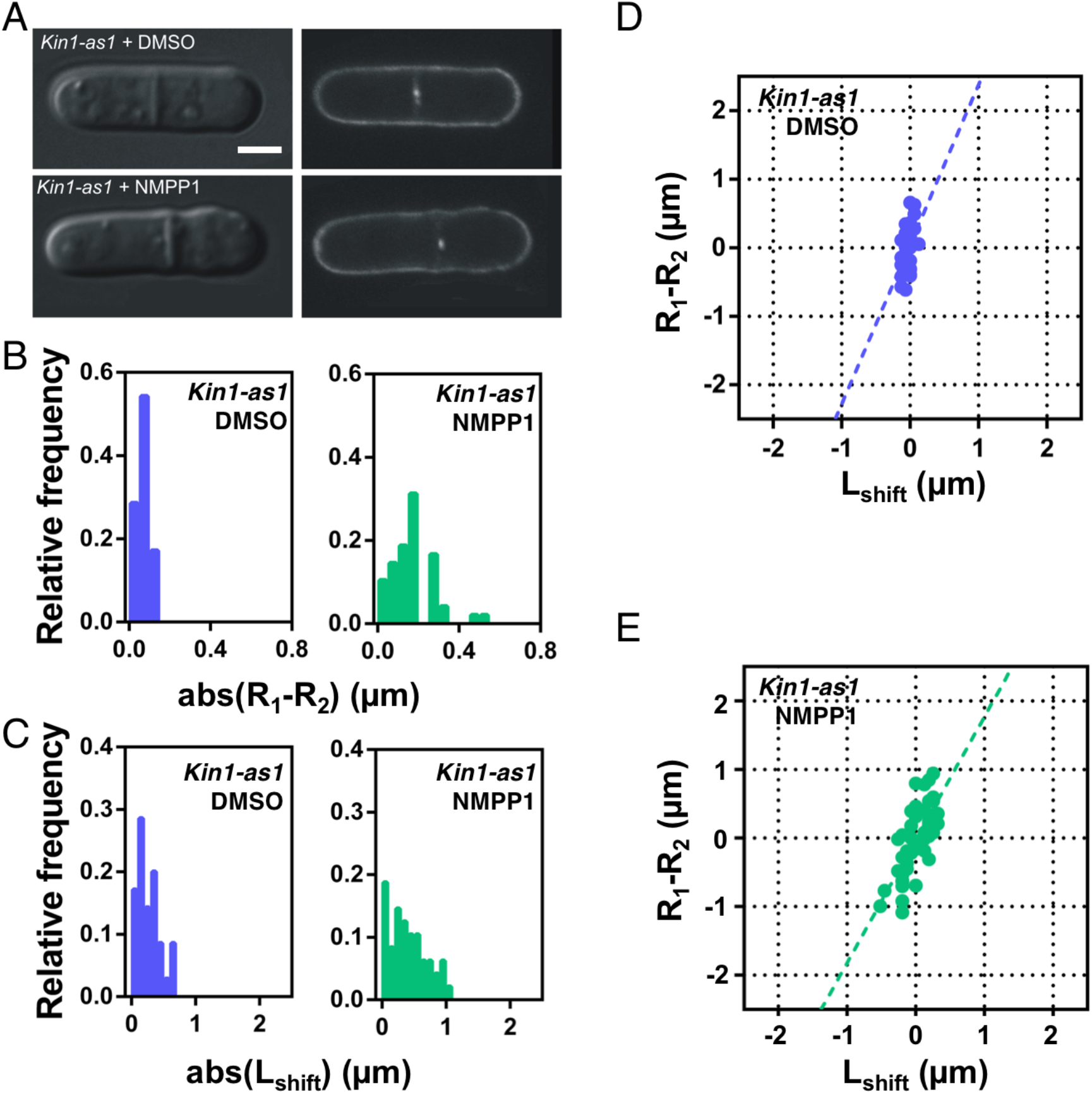
**(A)** DIC (left) and fluorescent (right) microscopy images (cdc15-GFP, isolectin-488) of kin1-as1 DMSO and kin1-as1 NMPP1 cells, Scale bar 2μm. **(B)** Distributions of the absolute values of Ro-R; for kin1-as1 DMSO and kin1-as1 NMPP1 cells. R_1_-R_2_ distributions are statistically different (Mann-Whitney, p < 0.0001). **(C)** Distributions of the absolute values of L_shift_ for kin1-as1 DMSO and kin1-as1 NMPP1 cells. Distributions are statistically different (t-test, p = 0.0122). **(D)** Correlation between L_shift_ and R_1_-R_2_ for kin1-as1 DMSO cells. The fit of Equation 1 is shown in the graph. **(E)** Correlation between L_shift_ and R_1_-R_2_ for kin1-as1 NMPP1 cells. The fit of Equation 1 is shown in the graph. n_Kin1-as1 DMSO_=35 and n_Kin1-as1 NMPP1_=48. Pearson’s correlation coefficient is 0.44 for kin1-as1 DMSO cells and Spearman correlation coefficient is 0.72 for kin1-as1 NMPP1 cells.

We monitored division site position and cell end curvatures using the method described above. *R_1_-R_2_* value is higher in *kin1-as1* NMPP1 cells (0.17±0.02 μm) compared to *kin1-as1* DMSO cells (0.06 ±0.01 μm), showing that cell end radii are less equivalent (Figure 2B). Furthermore, a larger *L_shift_* is clearly observed in *kin1-as1* NMPP1 cells (*L_shift_* = 0.57±0.04 μm) compared to *kin1-as1* DMSO cells (*L_shift_*=0.25±0.02 μm) (Figure 2C). This indicates that *kin1-as1* NMPP1 cells divide more asymmetrically than *kin1-as1* DMSO cells and WT cells. Again, *L_shift_*, plotted as a function of the *R_1_-R_2_* (Figure 2D and 2E), showed a positive correlation between *L_shift_* and *R_1_-R_2_* for *kin1-as1* NMPP1 cells compared to *kin1-as1* DMSO cells (correlation coefficient 0.72 and 0.44 respectively). *L_shift_* was again displaced towards the end with the highest radius of curvature, confirming the results with the *tea4Δ* mutant and strongly supporting the scaling laws of reference [16]. The mild correlation observed for *kin1-as1 DMSO* cells (0.44, Figure 2D) compared to the correlation seen for WT cells (0.03, Figure 1E) may reflect that the *kin1-as1* mutated allele is not fully functional in control condition as the wild type gene [24].

Applying Equation 1 and using the experimental values of L_shift_, R_2_, R_1_ and R (the radius of the cell at the middle plane) measured for *tea4Δ* cells and *kin1-as1* NMPP1 cells we could obtain the distribution of the mean wall surface tension <γ_end_> for each cell [17] (Figure 3). We then fitted the distributions with a gaussian function, obtaining a mean surface tension <γ_end_>= 4.9±3.1 N/m for *tea4Δ* cells and <γ_end_>= 2.9±3.4 N/m for *kin1-as1* NMPP1 cells, in good agreement in order of magnitude with values obtained with independent methods [17].

**Figure 3.**
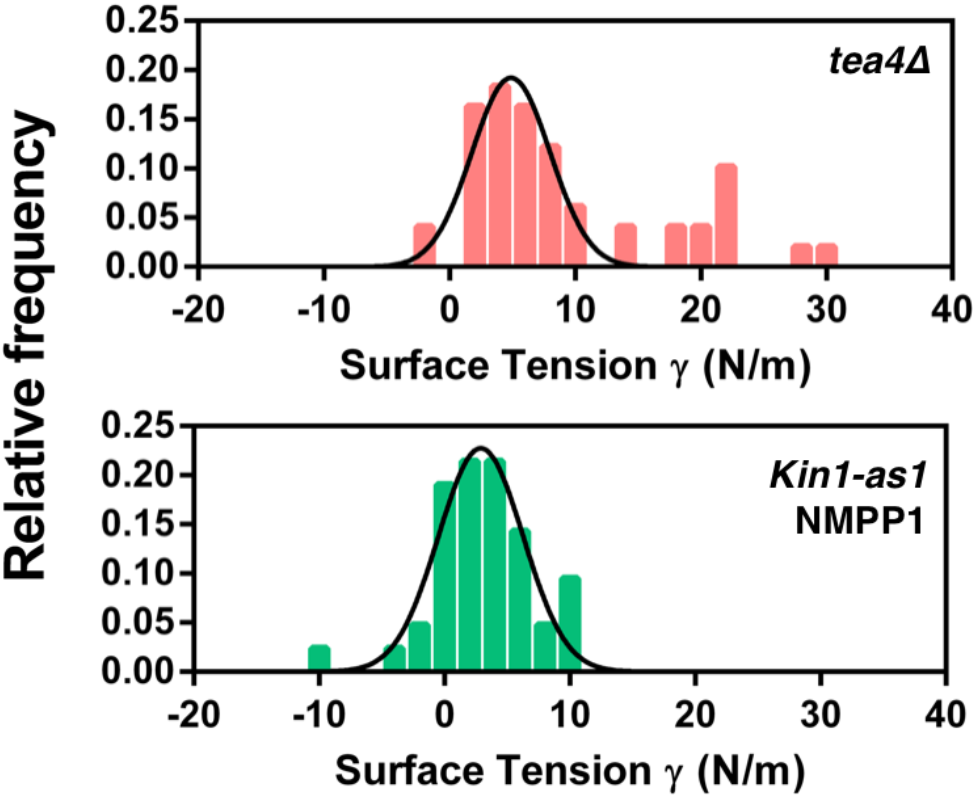
*Distribution of mean surface tension <γ> for tea4*Δ *cells and kin1-as1* NMPP1 *cells. <γ> was calculated according to Equation 1 assuming* Δ*P* = *0.85 MPa* [17]. *The data were fitted assuming a Gaussian distribution, obtaining a value for the surface tension of <γ_end_ >= 4.9±3.1 N/m for tea4Δ cells and <gend >= 2.9±3.4 N/m for kin1-as1* NMPP1 *cells*.

To further test the mechanism associated with shift in septum and curvatures at cell ends, we decided to measure wall thickness by performing measurements with Electron Microscopy ([25] and Figure 4A and 4B). The thickness was of the order of 200nm, as reported in other studies [27]. However, we systematically observed that one wall was thicker than the other (Figure 4C). This difference between the thick and the thin end was (45 ± 6 nm) and (47 ± 9 nm) for WT cells and *Kin1-as1 DMSO* cells respectively, but decreased by a 30% (32 ± 5 nm) when *Kin1-as1* cells were incubated with NMPP1. Finally, we found an inverse correlation between the cell end radius of curvature and wall thickness (Figure 4D). This is consistent with the notion that wall assembly is deficient in these mutants, and wall behaves as an inert ‘balloon’ with cylindrical shape similar to fission yeast: it undergoes thinning for larger radii of curvature.

**Figure 4.**
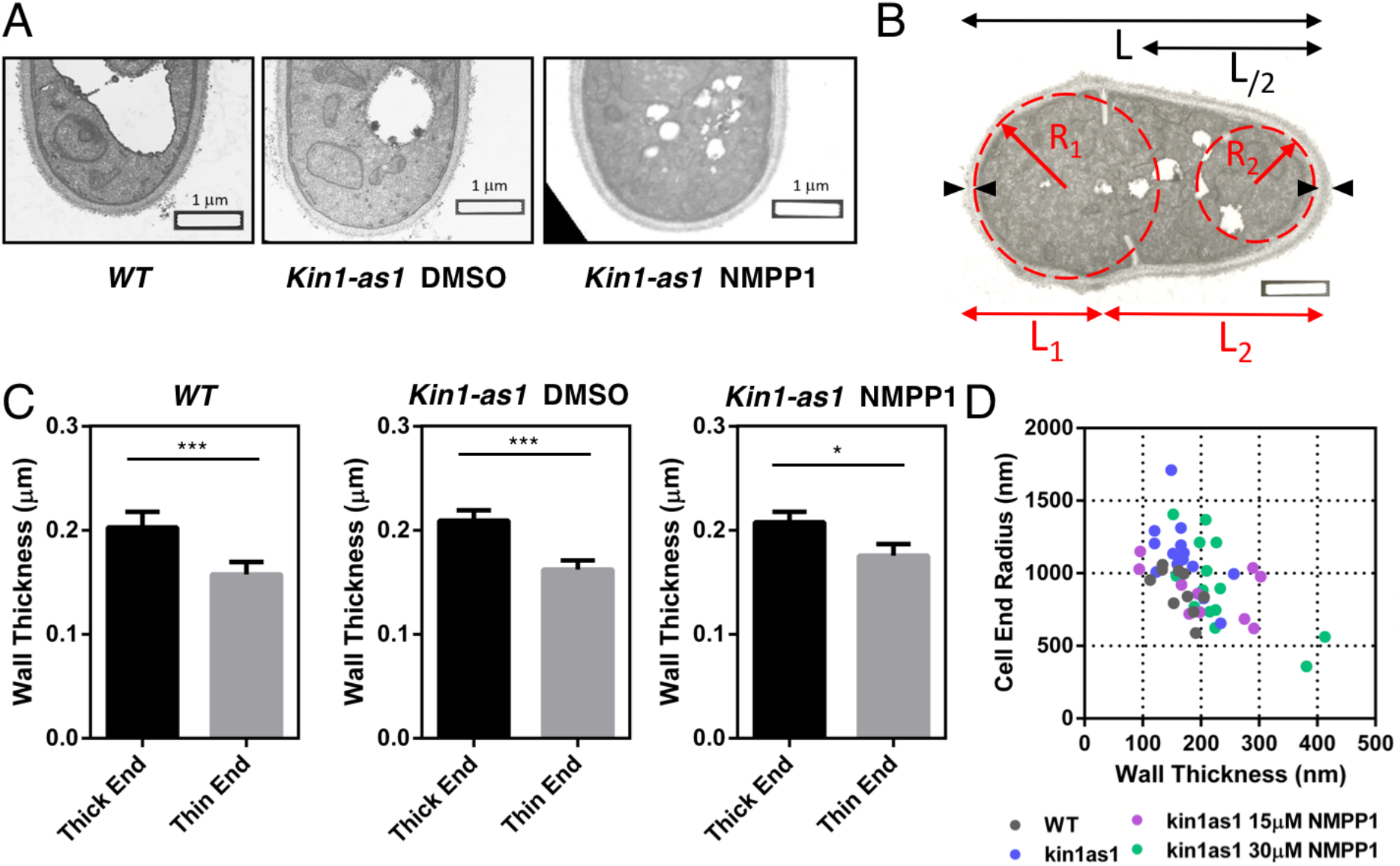
**(A)** Typical E.M. images for wild type, kin1-as1 and kin1-as1+NMPP1 cells. **(B)** Schematics of the parameters measured in each cell: R_i_ corresponds to the radii of curvature at the cell “end i”, and Li corresponds to the distance between “end i” and septum; L the total length and the wall thickness w is defined by the black arrows. **(C)** Measurements of the wall thickness. n_WT_=8, n_Kin1-as1 DMSO_=13 and n_Kin1-as1 NMPP1_=6. **(D)** Cell end radius as a function of wall thickness.

## 4. Discussion

Our study suggests an interplay between molecular actors of polarity, cell ends, and mechanics. The cell ends are defined by the concentration of polarity factors such as Tea1 and Bud6 that are absent from lateral sides. Tea1 acts to recruit other polarity factors, is itself delivered by microtubules and organizes the MT network, while Bud6 is known to bind to the F-actin nucleator For3 and regulates F-actin cable assembly and hence exocytic vesicle delivery [26,27]. In enlarged cell ends, these factors would tend to be less concentrated. The effect of increasing cell end radius on these aspects could be further studied through candidate mutant screens and imaging fluorescently tagged proteins. Cell end enlargement should also promote stretching of the cell wall, consistent with the thinning measured in mutants. This may activate mechanosensitive trans-membrane stress sensors such as Wsc1 and Mtl2 and/or the Cell Wall Integrity pathway [28,29]. Investigating their role by combining genetics, laser ablation, and soft-lithography techniques [30,31] would be valuable to link cell wall structure in contributing to the cell division control presented here.

Microtubules were shown to control the localization of the nucleus [18]. Both mechanisms could cooperate and compete. In this context, alternative mechanisms could be suggested. Microtubules may change their distributions and dynamics [32] in asymmetric cells. In turn, the localization of the nucleus could be shifted. Alternatively, microtubules dynamics *per se* could be altered by the curvatures at cell ends; in this case, the cell wall would act as the transmitter of asymmetry between ends directly. These alternative results could be tested with relevant mutants with modified microtubule dynamics and their live observation during septum formation.

The same approach could be used in other systems and for other biological functions, if we consider the cell cortex in other cells as the equivalent of cell wall in fission yeast [16]. For example, differentiation from stem cells has been associated to asymmetric cell division. In this context, the radii of curvatures of cells could be monitored over time and the onset of asymmetry may be associated as well with changes in radii of curvature at cell ends. To test this hypothesis, these experiments could be conducted for example in *Drosophila* [33,34], among other model systems. Similar approach may allow to give more quantitative substance to cortical force generators during cell division, in the context of *C. elegans* or mammalian cells [35]: this ‘gel’ localized at the cell poles spanning the cell membrane during cytokinesis may affect as well the localization of the cytokinetic furrow along the mechanism presented in this paper. In these different systems, future experiments could measure local curvatures at cell ends and cytokinesis localizations along our line to correlate poles shapes with respect to cytokinetic furrow plane. This could also be consistent with the notion that differences in tension between poles and furrow could play essential roles in cytokinesis [36]. Altogether ends and middles could be key read-outs in general for a variety of questions and model systems.

## 5. Conclusions

Our results show that fission yeast cell end shapes influence the division site position. In WT cells, the small difference in both cell end radii promotes balanced global forces that place the division site close to the geometric cell center. Accordingly, daughter cells divide at nearly equal sizes and this might be crucial for cell population fitness regarding symmetric partitioning of cellular components and damaged material inheritance [37]. We propose that two mechanisms contribute to symmetry of division in fission yeast: an ‘external’ input from cell wall driven forces and an ‘internal’ input driven by microtubule-dependent nuclear localization [18,19]. In mutants where the cell wall synthesis machinery is depolarized from cell ends but exhibit a normal microtubule network [22], the external cell wall contribution exceeds a threshold and cells divide asymmetrically, suggesting that the internal input cannot compensate the defect. The role of cell wall forces proposed here may be a generic mechanism in single celled symmetrically dividing organisms to produce equally sized daughter cells at each cell division.

## Acknowledgments

We thank A. Boudaoud (ENS Lyon) for his critical reading of the manuscript. This work was supported by a grant for “Aide aux financements de projets innovants interdisciplines” from the SFR BIOSIT (UMS CNRS 3480 - US INSERM 018, Rennes, France) to XLG, Unistra and CNRS funds to DR. This study was supported by the grant ANR-10-LABX-0030-INRT, a French State fund managed by the Agence Nationale de la Recherche under the frame program Investissements d’Avenir ANR-10-IDEX-0002-02. We thank J.R. Paulson and X. He (Oshkosh University, USA) for providing the NMPP1, A. Cadou and M. Sipiczki (University of Debrecen, Hungary) for TEM images, the MRic microscopy platform for fluorescence microscopy equipment at the SFR BIOSIT (Rennes, France), F. Chang (San Francisco, USA) and V. Simanis (Lausanne, Switzerland) for strains, J. Pécreaux (IGDR, Rennes, France) and A. Trubuil (INRA, Jouy-en-Josas, France) for their help in initial image analyses and Y. Arlot-Bonnemains (IGDR, Rennes, France) for support.

## References

1. S. L. Forsburg and others, Nat. Rev. Genet. 2, 659 (2001).

2. E. M. Jorgensen and S. E. Mango, Nat. Rev. Genet. 3, 356 (2002).

3. E. E. Patton and L. I. Zon, Nat. Rev. Genet. 2, 956 (2001).

4. D. St Johnston, Nat. Rev. Genet. 3, 176 (2002).

5. H.-O. Park, J. Hayles, C. Heichinger, K.-L. Hoe, L. Jeffery, D.-U. Kim, S. Salas-Pino, P. Nurse, and V. Wood, Open Biol. 3, 130053 (2013).

6. V. Graml, X. Studera, J. L. D. Lawson, A. Chessel, M. Geymonat, M. Bortfeld-Miller, T. Walter, L. Wagstaff, E. Piddini, and R. E. Carazo-Salas, Dev. Cell 31, 227 (2014).

7. F. Vaggi, J. Dodgson, A. Bajpai, A. Chessel, F. Jordán, M. Sato, R. E. Carazo-Salas, and A. Csikász-Nagy, PLoS Comput. Biol. 8, (2012).

8. O. Dudin, L. Merlini, F. O. Bendezú, R. Groux, V. Vincenzetti, and S. G. Martin, PLoS Genet. 13, e1006721 (2017).

9. N. Moris, J. Shrivastava, L. Jeffery, J.-J. Li, J. Hayles, and P. Nurse, Cell Cycle 15, 3121 (2016).

10. H. Cantwell and P. Nurse, PLoS Genet. 15, 1 (2019).

11. F. Chang, A. Woollard, and P. Nurse, J. Cell Sci. 109, 131 (1996).

12. C. Laplante, F. Huang, I. R. Tebbs, J. Bewersdorf, and T. D. Pollard, Proc. Natl. Acad. Sci. 113, E5876 (2016).

13. J. Dodgson, A. Chessel, M. Yamamoto, F. Vaggi, S. Cox, E. Rosten, D. Albrecht, M. Geymonat, A. Csikasz-Nagy, M. Sato, and R. E. Carazo-Salas, Nat. Commun. 4, 1834 (2013).

14. S. Hao, B. O’Shaughnessy, T. D. Pollard, D. Vavylonis, and J.-Q. Wu, Science. 319, 97 (2007).

15. A. Matsuyama, R. Arai, Y. Yashiroda, A. Shirai, A. Kamata, S. Sekido, Y. Kobayashi, A. Hashimoto, M. Hamamoto, Y. Hiraoka, S. Horinouchi, and M. Yoshida, Nat. Biotechnol. 24, 841 (2006).

16. D. Riveline, PLoS One 4, e6205 (2009).

17. N. Minc, A. Boudaoud, and F. Chang, Curr. Biol. 19, 1096 (2009).

18. P. T. Tran, L. Marsh, V. Doye, S. Inoué, and F. Chang, J. Cell Biol. 153, 397 (2001).

19. R. R. Daga and F. Chang, Proc. Natl. Acad. Sci. 102, 8228 (2005).

20. S. Moreno, A. Klar, and P. Nurse, Methods Enzymol. 194, 795 (1991).

21. H. Tatebe, K. Shimada, S. Uzawa, S. Morigasaki, and K. Shiozaki, Curr. Biol. 15, 1006 (2005).

22. S. G. Martin, W. H. McDonald, J. R. Yates, and F. Chang, Dev. Cell 8, 479 (2005).

23. A. Cadou, A. Couturier, C. Le Goff, T. Soto, I. Miklos, M. Sipiczki, L. Xie, J. R. Paulson, J. Cansado, and X. Le Goff, Mol. Microbiol. 77, 1186 (2010).

24. A. Cadou, A. Couturier, C. Le Goff, L. Xie, J. R. Paulson, and X. Le Goff, Biol. Cell 105, 129 (2013).

25. M. Osumi, Micron 29, 207 (1998).

26. J. M. Glynn, R. J. Lustig, A. Berlin, and F. Chang, Curr. Biol. 11, 836 (2001).

27. R. Behrens and P. Nurse, J. Cell Biol. 157, 783 (2002).

28. S. Cruz, S. Muñoz, E. Manjón, P. García, and Y. Sanchez, Microbiologyopen 2, 778 (2013).

29. P. P. and J. Cansado, Curr. Protein Pept. Sci. 11, 680 (2010).

30. C. R. Terenna, T. Makushok, G. Velve-Casquillas, D. Baigl, Y. Chen, M. Bornens, A. Paoletti, M. Piel, and P. T. Tran, Curr. Biol. 18, 1748 (2008).

31. V. Davì, L. Chevalier, H. Guo, H. Tanimoto, K. Barrett, E. Couturier, A. Boudaoud, and N. Minc, Proc. Natl. Acad. Sci. 116, 13833 (2019).

32. D. Foethke, T. Makushok, D. Brunner, and F. Nédélec, Mol. Syst. Biol. 5, 1 (2009).

33. S. Loubéry and M. González-Gaitán, Methods Enzymol. 534, 301 (2014).

34. E. Derivery, C. Seum, A. Daeden, S. Loubéry, L. Holtzer, F. Jülicher, and M. Gonzalez-Gaitan, Nature 528, 280 (2015).

35. S. Kotak and P. Gönczy, Curr. Opin. Cell Biol. 25, 741 (2013).

36. H. Turlier, B. Audoly, J. Prost, and J. F. Joanny, Biophys. J. 106, 114 (2014).

37. M. Coelho, S. J. Lade, S. Alberti, T. Gross, and I. M. Tolić, PLoS Biol. 12, 1 (2014).

